# Using joint models to adjust for informative drop-out when modelling a longitudinal biomarker: an application to type 2 diabetes disease progression

**DOI:** 10.1101/2021.06.17.448796

**Authors:** Harry D. Green, Katie G. Young, Angus G. Jones, Michael N. Weedon, John M. Dennis

## Abstract

Linear mixed effects models are frequently used in biomedical statistics to model the trajectory of a repeatedly measured longitudinal variable, such as a biomarker, over time. However, population-level estimates may be biased by censoring bias resulting from exit criteria that depend on the variable in question. A joint longitudinal-survival model, in which the exit criteria and longitudinal variable are modelled simultaneously, may address this bias. Using blood glucose progression (change in HbA1c) in type 2 diabetes patients on metformin monotherapy as an example, we study the potential benefit of using joint models to model trajectory of a biomarker in observational data. 7,712 patients with type 2 diabetes initiating metformin monotherapy were identified in UK Biobank’s general practice (GP) linked records. Genetic information was extracted from UK Biobank, and prescription records, baseline clinical features and biomarkers, and longitudinal HbA1c measures were extracted from GP records. Exit criteria for follow-up for a patient was defined as progression to an additional glucose-lowering drug (which is more likely in patient with higher HbA1c). Estimates of HbA1c trajectory over time were compared using linear mixed effect model approaches (which do not account for censoring bias) and joint models. In the primary analysis, a 0.19 mmol/mol per year higher (p = 0.01) HbA1c gradient was estimated using the joint model compared to the linear mixed effects model. This difference between models was attenuated (0.13 mmol/mol per year higher, p=0.43) when baseline clinical features and biomarkers were included as additional covariates.

Censoring bias should be carefully considered when modelling trajectories of repeatedly measured longitudinal variables in observational data. Joint longitudinal-survival models are a useful approach to identify and potentially correct for censoring bias when estimating population-level trajectories.

**Author Summary:** Modelling biomarkers that change over time in real world data is a challenging statistical problem due to many potential sources of bias. For example, when studying a chronic disease using a biomarker or other measurement that represents disease severity, medication intended to affect that measurement has a profound effect on how it will change over time. One common way to control for this is to study a cohort on the same treatment strategy. That way, results are not influenced by treatment change. If a patient progresses to stronger medication, then future data is no longer used. However, this approach introduces its own bias. Patients whose condition progress particularly quickly are more likely to change treatment more rapidly (and therefore be removed from further analysis, or ‘censored’), so the cohort is biased towards those whose condition progresses slower. In this paper we apply a technique called joint longitudinal-survival modelling which can adjust for this censoring bias and produce less biased estimates of progression rates. We use HbA1c (a widely used measure of glucose control) in type 2 diabetes as an example, however our methods are theoretically applicable to a range of problems across medicine in which a biomarker or feature is repeatedly measured in an individual.

## Introduction

Accurately modelling trajectories of a variable such as a clinical measure, biomarker or radiological measurement is of clinical interest across a range of biological and medical fields. For chronic, progressive diseases such as diabetes, inflammatory bowel disease, and prostate cancer, predicting the likely trajectory of a biomarker (HbA1c, faecal calprotectin and prostate-specific antigens respectively) is critical for understanding how the disease is likely to progress. However, if patients with more extreme trajectories are likely to be lost to follow-up, standard approaches to modelling longitudinal trajectories may be biased, as such patients contribute fewer data points to the model. This is particular problem in routine care based electronic health record datasets where protocol driven patient follow up is not available.

An example would be in modelling blood glucose control on a particular type 2 diabetes medication; patients with poor or rapidly deteriorating control may be more likely to have an event (e.g. loss to follow-up, death) or an intervention (e.g. initiation of another medication) that means follow-up data valid for the analysis become missing or censored at an earlier time point than patients with good glucose control. This informative drop-out could lead to the population level estimate being biased downwards, as the fastest progressors get censored quicker, contributing fewer data points to the model. Previous type 2 diabetes studies of blood glucose progression have not attempted to control for censoring bias [1,2].

We set out to explore the utility of joint longitudinal-survival modelling to estimate the trajectory of a biomarker in longitudinal population-based data, accounting for informative dropout [3,4]. It has previously been demonstrated in simulation study that joint longitudinal-survival modelling (JM) approach produces less-biased parameter estimates with more-reliable standard errors in the context of HIV viral and dynamic models [5]. Our case study is the modelling of long-term blood glucose control, as measured by HbA1c progression, in people with diabetes initiating metformin treatment.

## Methods

### Ethics Statement

Ethics approval for the UK Biobank study was obtained from the North West Centre for Research Ethics Committee (11/NW/0382) [6]. Written informed consent was obtained from all participants

### Data

The UK Biobank is a large-scale population-based study that aims to investigate the genetic and environmental basis of disease. Over 500,000 participants aged between 40 and 69 years were recruited between 2006 and 2010. Data collected include demographics, International Classification of Diseases 10th Revision (ICD10) hospital coding, medication records, anthropometric measures and a questionnaire containing lifestyle and mental-health factors. All participants have been genotyped, 438,427 using the Affymetrix Axiom UK Biobank array and 49,950 using the UK BiLEVE array. More detail on recruitment, demographics and data availability and the collection and imputation of genomic data can be found elsewhere [7].

Recently, the UK Biobank data on 230,105 participants have been linked with primary care record data [8]. These records are the only way participants’ health can be studied over time using the UK Biobank, and contain full information from a person’s longitudinal primary care records, including, critically for our study, routinely collected HbA1c records and drug prescription data.

### Study population

We considered people with type 2 diabetes on stable metformin monotherapy who were naïve to other glucose-lowering medication at the commencement of metformin. For each patient, the start of follow-up was from six months after initiation of metformin therapy, to exclude the period of acute drug response i.e. the fall in HbA1c commonly observed after initiating treatment, from the period of interest; the long term HbA1c progression on stable metformin therapy [9]. This approach is commonly used to model progression in diabetes [1,2,10].

In our study, patients were followed up until the earliest of: end of metformin prescription in the primary care record; initiation of another glucose-lowering medication; end of follow-up in the primary care record; 10 years after metformin initiation. Initiation of another glucose-lowering medication ending valid follow-up may lead to potentially informative drop-out in this study design; patients with faster progression (i.e. worsening HbA1c levels) will initiate additional treatment more quickly and so be lost to follow-up.

A full list of prescription codes used to define medication use is available in S1 Table. Patients without an HbA1c record within the study period were excluded.

### Baseline clinical features and biomarkers

Clinical features and biomarkers (full list below) were extracted from the primary care record at the time of metformin initiation (baseline), using previously described procedures [11]. While for the longitudinal modelling, HbA1c values were only considered valid 6 months after the onset of metformin, baseline variables were considered to be the closest recording to the date of metformin initiation, within a 6-month window either side. We considered the following baseline clinical features and biomarkers, which have previously been demonstrated to associate with diabetes progression; HbA1c, age at starting metformin, sex, weight, HDL-cholesterol (HDL-c), Triglycerides, type 1 diabetes genetic risk score as defined in [12], and duration of type 2 diabetes[13].

Type 2 Diabetes was defined in primary care data as any two of: a diagnosis code for diabetes (see S2 Table), an HbA1c measurement ≥ 48 mmol/mol, and a prescription for glucose-lowering medication in the primary care record, where diagnosis date is defined as the earliest occurrence. Diabetes duration was defined as the time period between diagnosis and metformin initiation. For some individuals, the earliest evidence of diabetes was metformin initiation, resulting in a large amount of 0 values for diabetes duration. As a result, this was split into categories 0-1, 1-2, 2-5, 5-10, 10+ years of diabetes prior to commencing metformin for statistical analyses.

### Study Outcomes

We jointly modelled glycemic progression, as a longitudinal repeatedly measured outcome, and treatment progression, the initiation of another glucose-lowering medication, as a time-to-event outcome. All HbA1c recordings during the study period (from 6 months to 10 years after metformin initiation) were included as longitudinal outcomes.

### Statistical Methods

We compared linear estimates of HbA1c progression over time estimated from two modelling approaches: a ‘standard’ linear mixed effects (LME) model (incorporating solely the longitudinal HbA1c outcome) and a joint longitudinal-survival model (JM) (incorporating both the longitudinal HbA1c outcome and the treatment progression time to event outcome). An underlying random effects model is used to link the longitudinal and survival outcomes[3,4]. We used the R package ‘JM’ for all JMs in this paper and conducted analysis in R 3.5.2 [14].

Initial models adjusted for baseline HbA1c and age at metformin treatment initiation, features that were available for all patients and those most strongly associated with progression in previous research [15,16].

Secondly, we updated the initial models to include the additional clinical features. We tested a model additionally adjusting for weight, HDL-c and log triglycerides (TG). We also included sex, duration of diabetes, and genetic risk scores for type 1 diabetes. The purpose of this analysis was to test whether including additional covariates in a LME model can address any potential bias, and also whether the censoring bias affects observed associations between clinical covariates and HbA1c progression.

As a sensitivity analysis we refitted each model to include all HbA1c records from the start of metformin (rather than from 6 months after initiation). To allow for the initial period of response we fitted a linear spline that with a breakpoint at 6 months to allow for nonlinearity of the HbA1c trajectory.

Patients with missing data were excluded. We did not consider multiple imputation, as these readings are taken in primary care, and missingness will not be at random[17]. Descriptive summary statistics were calculated using the cohort with complete clinical feature and biomarker information, excluding those with missing data.

A p-value was derived to test for differences between model estimates using a t test, where the z-score was calculated by 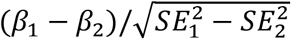.

## Results

7,764 people initiating metformin were eligible for the study as their first-line glucose lowering medication had a period on unchanged treatment longer than 6 months, a valid baseline HbA1c record, and at least one valid HbA1c recorded at least 6 months after initiating treatment (i.e. in the ‘progression’ period) (Figure 1). 7,712 (99.34%) had evidence of type 2 diabetes in either the UK Biobank self-report, HES, or primary care data, and this sample was used for the simple model.

**Figure 1.**
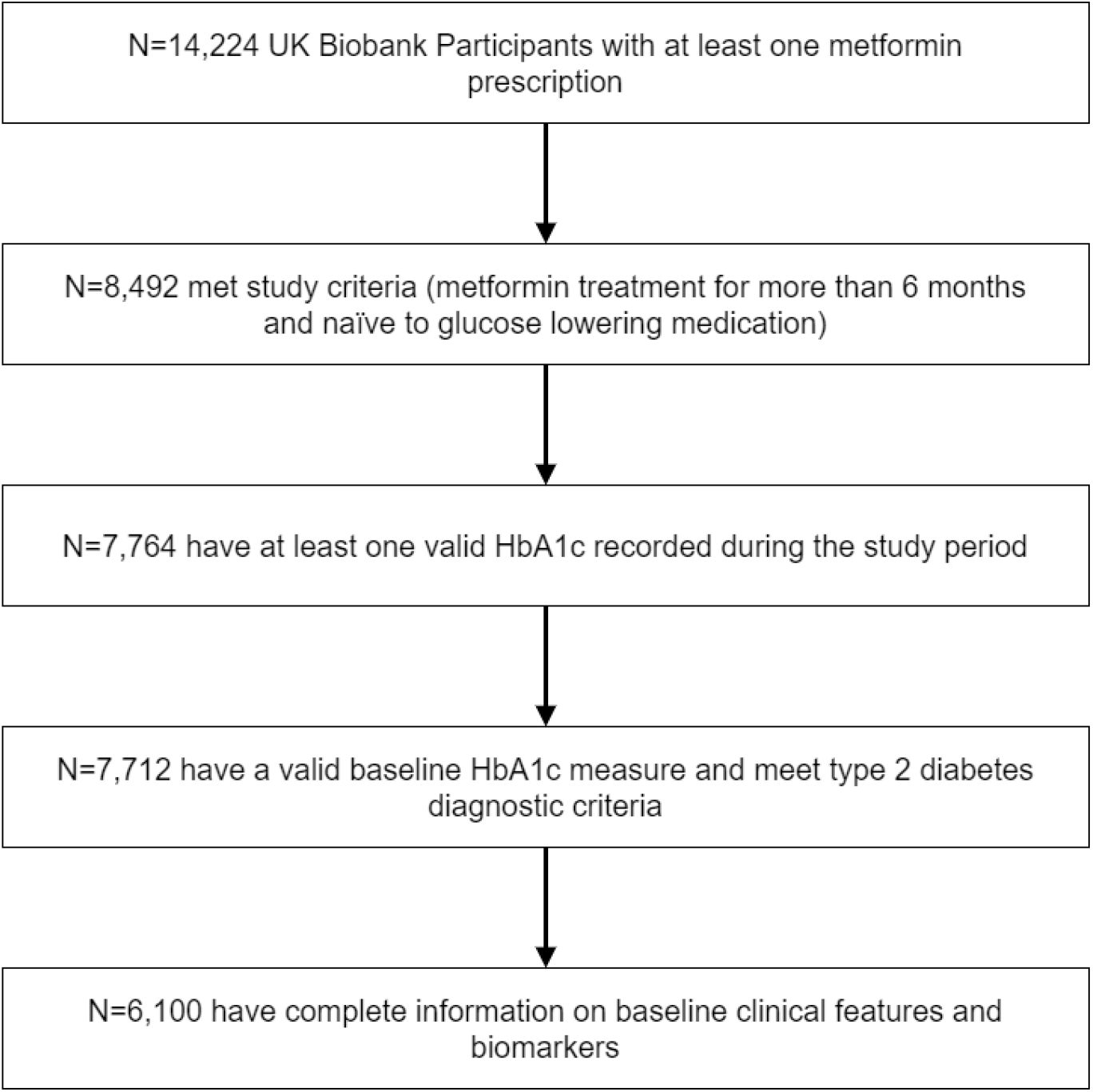
A flowchart showing numbers in the model.

Mean follow-up time was 3.1 (SD 2.0) years in those who progressed to additional therapy and 4.1 (SD 2.7) in those who did not progress. A Kaplan-Meier curve of cumulative incidence of progression to additional therapy can be found in Supplementary Figure 1. During this period, participants provided a mean of 6.5 HbA1c recordings (SD 5.1). Estimated 10 year incidence of progression to additional therapy was 43.26% (41.00-45.45).

Patients who progressed to second-line therapy tended to be younger, and had a higher weight, initial HbA1c and log triglycerides, a longer duration of diabetes, and had a lower HDL than those that did not progress (Table 1). There was no significant difference in progression status by sex or genetic risk of T1D.

**Table 1.**

| Variable (unit) | Progress<br>(N=1,221) | No progress<br>(N=4879) | p-value |
| --- | --- | --- | --- |
| Starting Age (years) | 58.58 +/- 7.67 | 61.50 +/- 7.57 | $1 \times 10^{-32}$ |
| Initial Weight (kg) | 94.85 +/- 19.01 | 91.46 +/- 18.65 | $3 \times 10^{-8}$ |
| Initial HbA1c (mmol/mol) | 68.38 +/- 18.64 | 61.75 +/- 16.87 | $2 \times 10^{-30}$ |
| T1D GRS | 10.30 +/- 2.50 | 10.16 +/- 2.42 | 0.055 |
| HDL (mmol/L) | 1.17 +/- 0.30 | 1.21 +/- 0.32 | $9 \times 10^{-5}$ |
| log TG (mmol/L) | 0.65 +/- 0.50 | 0.56 +/- 0.49 | $3 \times 10^{-9}$ |
| Diabetes Duration (years) | 3.44 +/- 7.68 | 2.82 +/- 5.43 | $8 \times 10^{-6}$ |
| Male sex % (N) | 63.39% (774) | 60.89% (2971) | 0.13 |

Baseline patient characteristics and biomarkers, split by treatment progression status

6,100 people (79.10% of those in cohort 1) had all baseline clinical features recorded. This complete case cohort were significantly older: the mean age in the 6,100 with all data available was 60.9 compared to 58.1 in the 1,663 excluded due to missingness (p=1&10^−37^). A table showing the distributions of the covariates used in the 6,100 cohort is given in table 1, split by whether or not the patient progressed to second-line medication.

### Estimates of HbA1c progression are greater using a joint model than a standard approach

The estimated rate of HbA1c progression was greater using the JM (incorporating informative drop-out) then the linear mixed model (Figure 2). Using the JM, the estimated rate of progression was 2.10 mmol/mol per year (95% confidence interval 2.00, 2.20) compared to 1.91 (95% confidence interval 1.81, 2.02) in the LME model (p value of 0.01 for test of difference in gradient). Full longitudinal fixed effect estimates for each model can be found in S3 Table.

**Figure 2.**
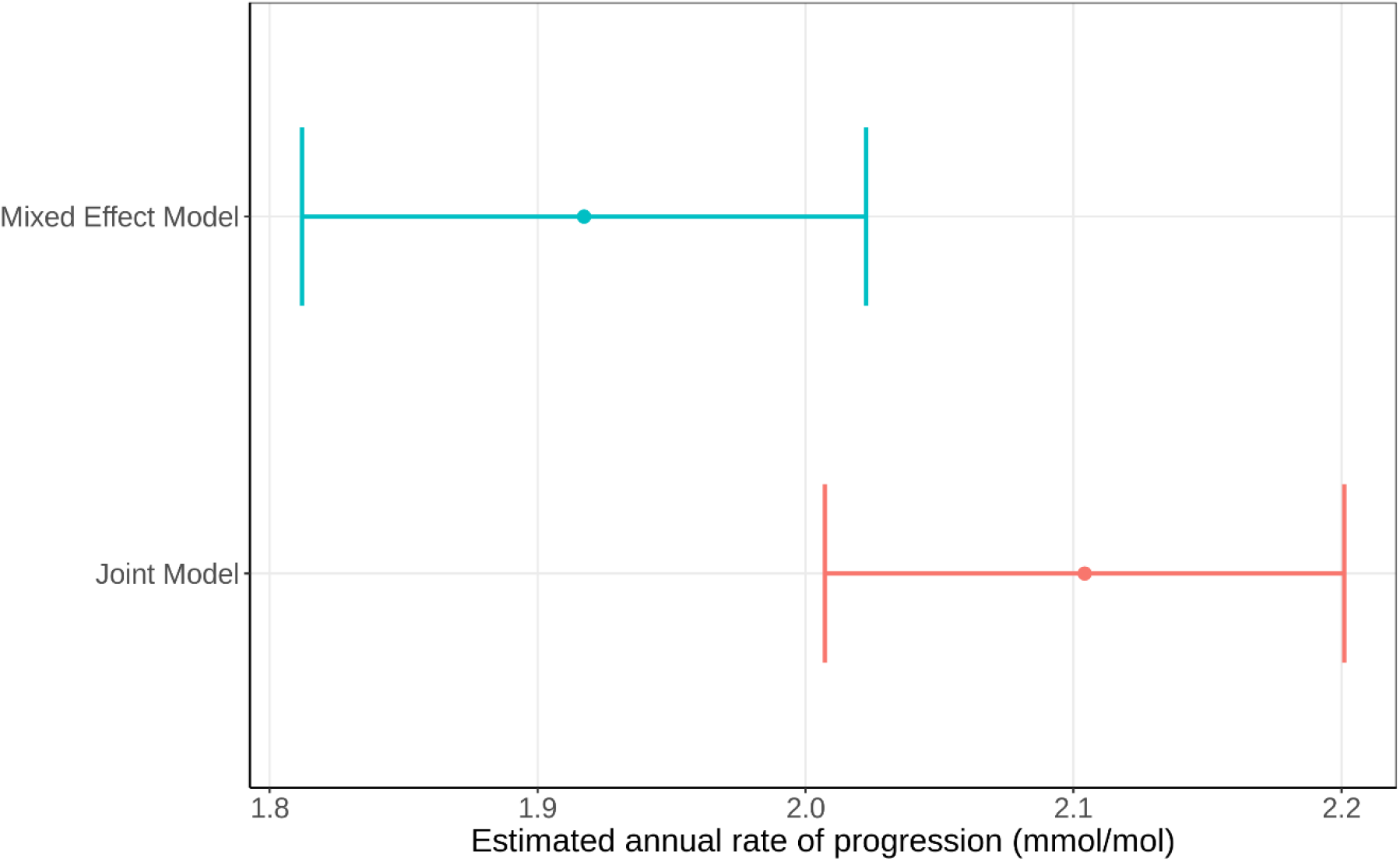
Estimated rate of type 2 diabetes progression (annual rate of deterioration in HbA1c in mmol/mol) using the simple joint model and the simple linear mixed-effects model (n=7,712)

When including the initial 6 months of HbA1c measures using a linear spline, the population level gradient for the post 6 month period using the JM was near identical to the primary analysis (2.1 mmol/mol per year (95% confidence interval 2.0, 2.2).

### Differences between the joint model and standard model are attenuated when additional clinical variables are added

The difference between the JM and the LME model was attenuated when more explanatory variables were used. Fitting a multivariable model adjusting for the full covariate set (S4 Table), the JM returned a gradient of 1.92 (1.70, 2.14), and the LME model: 1.79 (1.57,2.02) [p=0.42 for difference in gradient]. Estimates for the associations between each clinical covariate and the rate of progression were near identical using the JM and LME approaches (Figure 3). Full longitudinal fixed effect estimates for each model can be found in S4 Table.

**Figure 3.**
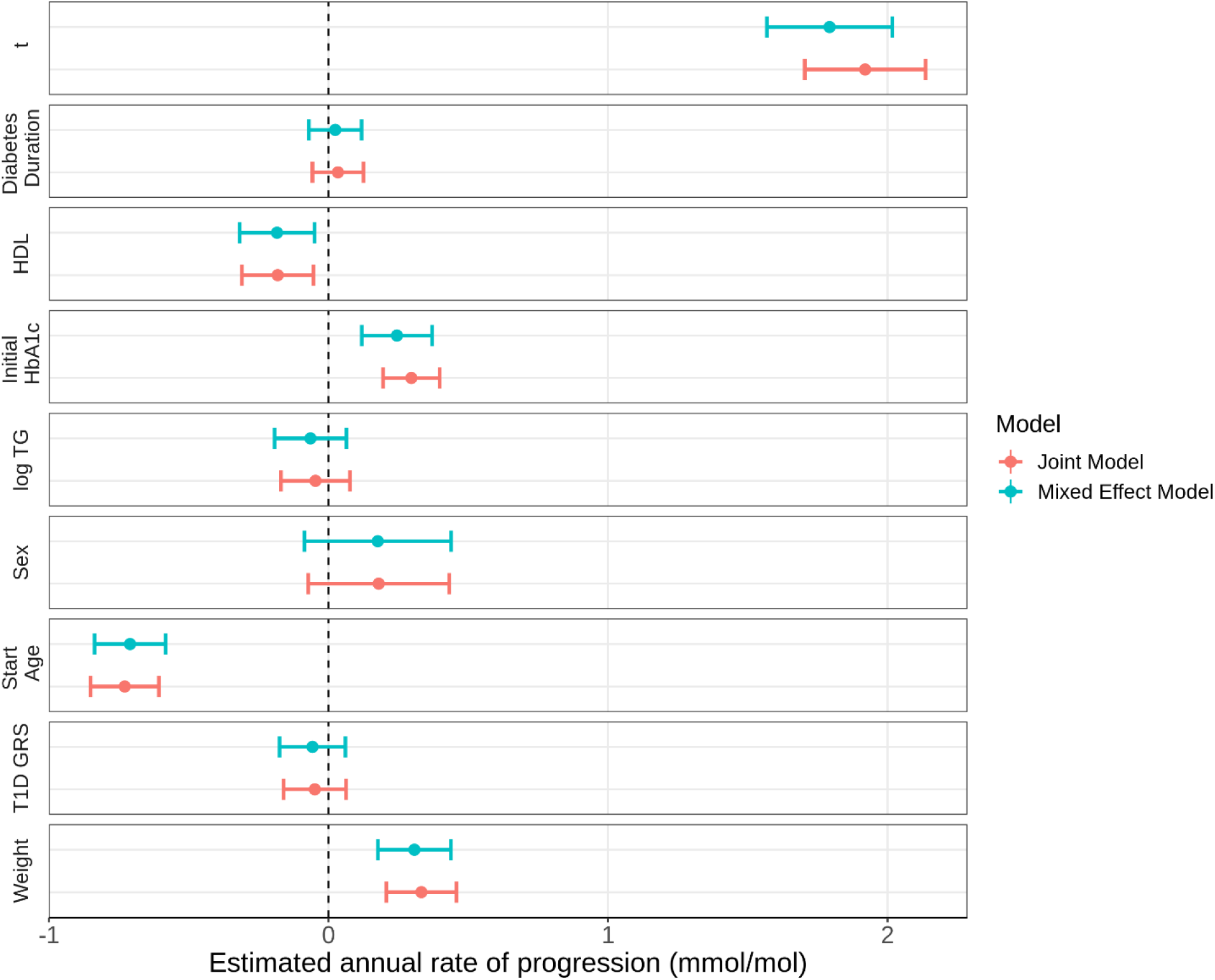
Estimates of the effect of clinical covariates on rate of diabetes progression for both the joint model and the linear mixed-effects model. t represents the average rate of HbA1c progression. Other variables’ estimates represent their effect on the rate of progression (N=6,100).

## Discussion

### Principal Findings

We have demonstrated that the rate of biomarker progression estimated from population-based observational data is confounded if censoring bias is not incorporated into the underlying estimation model. A simple JM including starting HbA1c and age as covariates, returned a significantly different estimate to a standard LME model for the population level HbA1c gradient. When adjusting for clinical features previously demonstrated to be associated with diabetes progression, the difference between the models was attenuated and not statistically significant with the available sample size. This highlights the importance of informed covariate selection when modelling specific biomarker trajectories.

A further important finding, relevant for studies focused on precision medicine (i.e. discovery of individual-level features modifying trajectory of a biomarker) is that estimated effect sizes for associations between clinical covariates and rate of HbA1c progression were stable between the complex JM and the simpler LME model. This, however, should be considered a finding specific to our study; it is possible that in other analyses informative dropout might affect such estimates. Future simulation studies to evaluate the potential benefit from applying JM’s in a precision medicine context would be of considerable interest.

### Strengths and Weaknesses

This was the first study to use a JM to study diabetes progression and the first to investigate diabetes progression using the UK Biobank’s GP records. This gives us the advantage of using real-world data to assess the population-level HbA1c gradient in a real-world setting. The use of the UK Biobank allows us to study genetics alongside this data, and we were able to include genetic risk of both type 1 diabetes in a clinical model.

Limitations of the study result from the nature of primary care data. As this data was recorded in clinic, time-dependent variables, such as HbA1c, are not recorded at regular intervals, and temporal resolution of these records vary between patients. While the clinical covariates such as weight, triglycerides and HDL were recorded in everyone at the UK Biobank assessment centre, this visit would have occurred at different times in disease course. As a result, we used GP records for these clinical covariates close to the start date, which are not available for every patient, which cannot be imputed as primary care missingness is likely to be not at random (for example, patients with greater comorbidity are likely to have standard blood tests more frequently than healthier patients). Further, while we consider the first record of a metformin prescription to correspond to the beginning of therapy, it is possible these individuals had a prescription to metformin, or another glucose-lowering drug, prior to the recorded date in the GP records. It is important to note the computational intensity of using a JM over a LME model. The simple JM model took 12 times longer than the LME counterpart, and 6 times longer when additional covariates were used.

### Implications

Our results demonstrate that one must be bias from informative drop-out should be carefully considered when attempting to model longitudinal trajectories in observational data. We propose that the JM approach should be used in such situations in preference of LME models, or at least could serve as a useful sensitivity analysis to assess degree of censoring bias in future studies. Although our case study was specific to type 2 diabetes, our findings have implications for the study of trajectories of a longitudinal variable across many common diseases.

## Acknowledgements

This research has been conducted using the UK Biobank Resource (Application number: 9055). The authors acknowledge the use of the University of Exeter High-Performance Computing facility in carrying out this work.

## Supporting Information Legend

**S1_Table.xlsx**

S1 table - A list of glucose-lowering medication drugs used in the paper

**S2_Table.xlsx**

S2 table - A list of read codes for diabetes diagnosis used in the paper

**S3_Table.docx**

S3 table - Longitudinal effects in the Joint Model and the Linear Mixed Effects Model

**S4 Table.docx**

Longitudinal effects in the Joint Model and the Linear Mixed Effects Model with covariates

**S1_Fig.tif**

Kaplan Meier curve of treatment progression

## References

1. Goswami S, Yee SW, Xu F, Sridhar SB, Mosley JD, Takahashi A, et al. A Longitudinal HbA1c Model Elucidates Genes Linked to Disease Progression on Metformin. Clin Pharmacol Ther. 2016;100: 537–547. doi:10.1002/cpt.428

2. Chalmers J, Hunter JE, Robertson SJ, Baird J, Martin M, Franks CI, et al. Effects of early use of pioglitazone in combination with metformin in patients with newly diagnosed type 2 diabetes. Curr Med Res Opin. 2007;23: 1775–1781. doi:10.1185/030079907X210606

3. Sudell M, Kolamunnage-Dona R, Tudur-Smith C. Joint models for longitudinal and time-to-event data: a review of reporting quality with a view to meta-analysis. BMC Med Res Methodol. 2016;16: 168. doi:10.1186/s12874-016-0272-6

4. Rizopoulos D. Joint models for longitudinal and time-to-event data: With applications in R. CRC press; 2012.

5. Wu L. A Joint Model for Nonlinear Mixed-Effects Models With Censoring and Covariates Measured With Error, With Application to AIDS Studies. J Am Stat Assoc. 2002;97: 955–964. doi:10.1198/016214502388618744

6. Bycroft C, Freeman C, Petkova D, Band G, Elliott LT, Sharp K, et al. The UK Biobank resource with deep phenotyping and genomic data. Nature. 2018;562: 203–209. doi:10.1038/s41586-018-0579-z

7. Sudlow C, Gallacher J, Allen N, Beral V, Burton P, Danesh J, et al. UK biobank: an open access resource for identifying the causes of a wide range of complex diseases of middle and old age. PLoS Med. 2015;12: e1001779. doi:10.1371/journal.pmed.1001779

8. UK Biobank. Primary Care Linked Data :Companion Document. 2019; 1–24. Available: https://www.ukbiobank.ac.uk

9. Kahn SE, Haffner SM, Heise MA, Herman WH, Holman RR, Jones NP, et al. Glycemic Durability of Rosiglitazone, Metformin, or Glyburide Monotherapy. N Engl J Med. 2006;355: 2427–2443. doi:10.1056/NEJMoa066224

10. Donnelly LA, Zhou K, Doney ASF, Jennison C, Franks PW, Pearson ER. Rates of glycaemic deterioration in a real-world population with type 2 diabetes. Diabetologia. 2017/12/19. 2018;61: 607–615. doi:10.1007/s00125-017-4519-5

11. Rodgers LR, Weedon MN, Henley WE, Hattersley AT, Shields BM. Cohort profile for the MASTERMIND study: Using the Clinical Practice Research Datalink (CPRD) to investigate stratification of response to treatment in patients with type 2 diabetes. BMJ Open. 2017;7. doi:10.1136/bmjopen-2017-017989

12. Sharp SA, Rich SS, Wood AR, Jones SE, Beaumont RN, Harrison JW, et al. Development and Standardization of an Improved Type 1 Diabetes Genetic Risk Score for Use in Newborn Screening and Incident Diagnosis. Diabetes Care. 2019; dc181785. doi:10.2337/dc18-1785

13. Zhou K, Donnelly LA, Morris AD, Franks PW, Jennison C, Palmer CNA, et al. Clinical and Genetic Determinants of Progression of Type 2 Diabetes: A DIRECT Study. Diabetes Care. 2014;37: 718LP– 724. doi:10.2337/dc13-1995

14. Rizopoulos D. JM: An R Package for the Joint Modelling of Longitudinal and Time-to-Event Data. J Stat Softw. 2010;35. doi:10.18637/jss.v035.i09

15. Zhou K, Donnelly LA, Morris AD, Franks PW, Jennison C, Palmer CNA, et al. Clinical and genetic determinants of progression of type 2 diabetes: A DIRECT Study. Diabetes Care. 2013; DC_131995. doi:10.2337/dc13-1995

16. Dennis JM, Shields BM, Henley WE, Jones AG, Hattersley AT. Disease progression and treatment response in data-driven subgroups of type 2 diabetes compared with models based on simple clinical features: an analysis using clinical trial data. lancet Diabetes Endocrinol. 2019;7: 442–451. doi:10.1016/S2213-8587(19)30087-7

17. Marston L, Carpenter JR, Walters KR, Morris RW, Nazareth I, Petersen I. Issues in multiple imputation of missing data for large general practice clinical databases. Pharmacoepidemiol Drug Saf. 2010;19: 618–626. doi:10.1002/pds.1934

