## Supplementary figures and images for "Using joint models to adjust for informative drop-out when modelling a longitudinal biomarker: an application to type 2 diabetes disease progression"

### Supplementary Figure 1

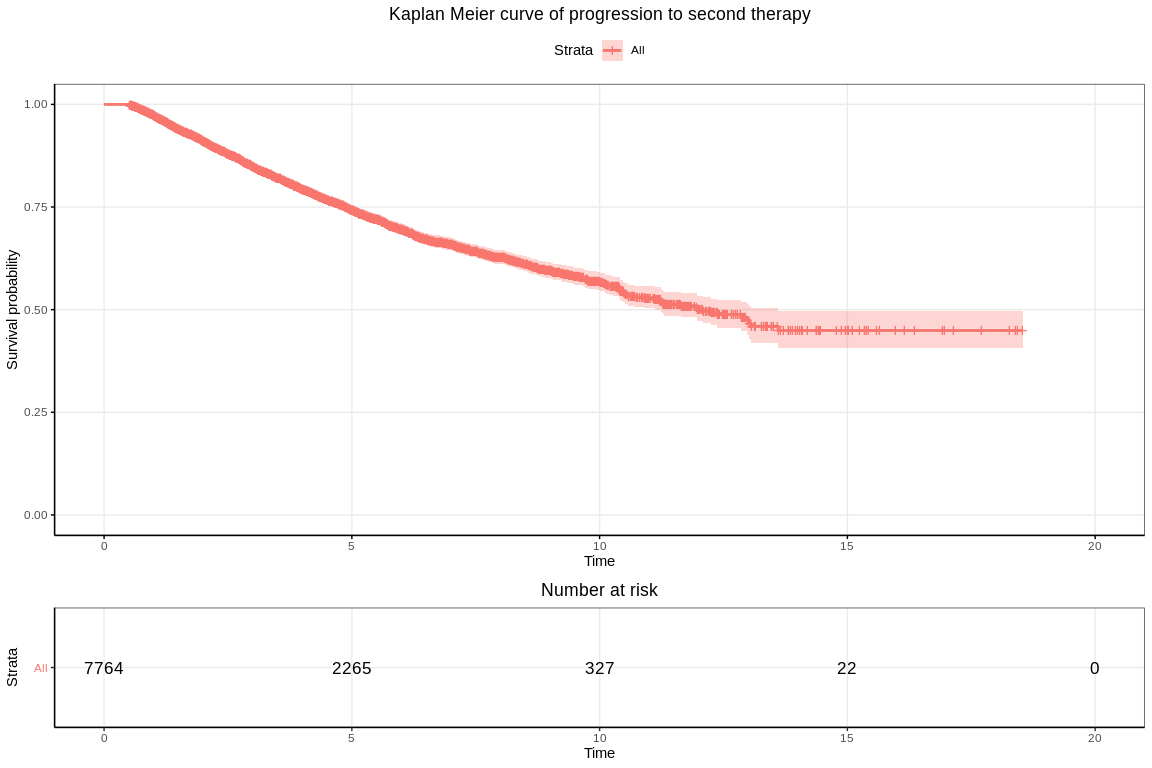
