## Supplementary material for "Using joint models to adjust for informative drop-out when modelling a longitudinal biomarker: an application to type 2 diabetes disease progression": S3 Table

### Supplementary Table 1: Fixed Effects of the Simple Model

Longitudinal effects in the Joint Model and the Linear Mixed Effects Model

|  | JM Estimate | JM SE | JM p-value | LME Estimate | LME SE | LME p-value |
| --- | --- | --- | --- | --- | --- | --- |
| (Intercept) | 51.61 | 0.13 | <10^-300^ | 51.87 | 0.13 | <10^-300^ |
| Time (years) | 2.10 | 0.05 | <10^-300^ | 1.92 | 0.05 | 8x10^-275^ |
| Start Age (years) | 0.02 | 0.02 | 0.28 | 0.01 | 0.02 | 0.69 |
| Initial HbA1c (mmol/mol) | 0.22 | 0.01 | 1x10^-189^ | 0.22 | 0.01 | 2x10^-173^ |
| t:Start Age | -0.10 | 0.01 | 2x10^-65^ | -0.10 | 0.01 | 2x10^-65^ |
| t:Initial HbA1c | 0.01 | 0.00 | 1x10^-8^ | 0.01 | 0.00 | 3x10^-3^ |
