## Supplementary material for "Using joint models to adjust for informative drop-out when modelling a longitudinal biomarker: an application to type 2 diabetes disease progression": S4 Table

### Supplementary Table 2: Fixed Effects of the Model with Clinical Covariates

Longitudinal effects in the Joint Model and the Linear Mixed Effects Model with covariates

|  | JM Estimate | JM SE | JM p-value | LME Estimate | LME SE | LM p-value |
| --- | --- | --- | --- | --- | --- | --- |
| t | 1.92 | 0.11 | <10^-300^ | 1.79 | 0.11 | <10^-300^ |
| t:Start Age | -0.73 | 0.06 | 1x10^-31^ | -0.71 | 0.06 | 8x10^-28^ |
| t:Initial HbA1c | 0.30 | 0.05 | 1x10^-8^ | 0.24 | 0.06 | 1x10^-4^ |
| t:T1D GRS | -0.05 | 0.06 | 0.39 | -0.06 | 0.06 | 0.34 |
| t:HDL | -0.18 | 0.07 | 5x10^-3^ | -0.18 | 0.07 | 7x10^-3^ |
| t:TG | -0.05 | 0.06 | 0.46 | -0.06 | 0.07 | 0.33 |
| t:Weight | 0.33 | 0.06 | 2x10^-7^ | 0.31 | 0.07 | 4x10^-6^ |
| t:Sex | 0.18 | 0.13 | 0.16 | 0.18 | 0.13 | 0.19 |
| t: Diabetes Duration | 0.03 | 0.05 | 0.47 | 0.02 | 0.05 | 0.62 |
